# Diagnostic of begomoviruses in complex infections –a case study

**DOI:** 10.1101/2020.01.19.911776

**Authors:** Juan Jovel

**Affiliations:** Pacific Agriculture and Agri-food Canada. 4200 Highway 97, Summerland, BC. VOH 1ZO

**Keywords:** Begomoviruses, diagnostic, epidemics, mix infections, recombination

## Abstract

Complexes of viruses inducing a syndrome in plants strongly hinder the identification of the causal agent of a disease. The *Sida micrantha* mosaic disease is associated with a complex of begomoviruses. For more than twenty years, two DNAs A (DNA A2 and DNA A3) belonging to this complex could neither be detected nor isolated from *Sida micrantha* Schr. plants, although one of them (DNA A2) now appears to be the major component of the complex. A random unintended *Bemisia tabaci*-mediated transmission of begomoviruses from several Sida species – including *S. micrantha* – to experimental *Malva parviflora* plants resulted in the serendipitous finding of these new DNAs A. Simultaneously, a number of other begomoviruses infecting Sida plants from several Latin American countries were transmitted to *M. parviflora* plants and the convergence of them resulted in natural pseudorecombinants. Pseudorecombination, however, took place exclusively between heterologous genomic components that shared identical binding sites for the replication-associated protein AC1. This case study constitutes an exceptional opportunity for the analysis of plant–virus– vector interactions in a quasi-natural environment. In addition to that, the methodology described here may be used to isolate and characterize different begomoviruses inducing a syndrome during mix infections.

## Introduction

In several aspects, geminiviruses (Rybicki *et al.*, 2000) are intriguing pathogens of plants. From an economical perspective they are responsible for extensive losses in diverse cash crops. For instance, African cassava mosaic (Thottappilly, 1992), tomato leaf curl (Czosnek and Laterrot, 1997), cotton leaf curl (Briddon and Markham, 2001), and bean golden mosaic (Morales and Anderson, 2001) diseases have been major bottlenecks in the production of vegetables and fibber crops worldwide during the last three decades. Because of their limited coding capacity, geminiviruses strongly rely on host replication complexes to accomplish their life cycle. This peculiarity makes them interesting models for the study of replication, transcription and cell cycle regulation in higher plants (Hanley-Bowdoin *et al.*, 1999). Replication of geminiviruses is assumed to occur through a rolling circle mechanism (RCR) (Hanley-Bowdoin *et al.*, 1999) complemented by a recombination-dependent (RDR) mechanism (Jeske *et al.*, 2001).

The *Geminiviridae* family comprises four genera: *Mastreviruses*, *Begomoviruses*, *Topocuviruses*, and *Curtoviruses*. While begomoviruses may possess mono– or bipartite genomes, all members of the other groups are strictly monopartite. In bipartite begomoviruses, the genomic components are designated DNA A and DNA B and possess divergent nucleotide sequence with the exception of a ~200 nucleotide-long common region (CR), harbouring the origin of replication and promoters. DNA A codes for the proteins associated with the viral replication (AC1 and AC3), transcriptional regulation (AC1 and AC2), and the coat protein (AV1), whereas DNA B codes for a nuclear shuttle (BV1) and a cell-to-cell movement proteins (BC1) (Rybicki *et al.*, 2000). Since DNA B does not possess a replication-associated protein, its replication depends on the DNA A-encoded AC1 protein and takes place in a virus-specific manner. While all determinants of such specificity are not known at the moment, it has been demonstrated that a direct repeat located in the CR and matching the consensus sequence 5’–**GG–A**GTAYY**GG–AG** is necessary, although not sufficient, to confer specific AC1 binding to DNA B (Hanley-Bowdoin *et al.*, 1999). Thus, the existence of a DNA B component implies the presence of its cognate DNA A.

*Sida micrantha* mosaic disease is associated with a complex of begomoviruses that have been partially characterized (Jovel *et al.*, 2004; Jovel *et al.*, 2007). Unraveling the first pieces of this puzzle has been a 25-years-long effort. Initially, a DNA A (DNA A1) and a DNA B (DNA B1) were cloned and sequenced. The weak homology between DNA A1 and DNA B1 in the common region (CR) suggested that they were components of different begomoviruses. Almost a decade later, two additional DNAs B (DNA B2 and DNA B3) were cloned directly from the Sida plants and sequenced. The sequences of the CR of DNA B3 and DNA A1 were highly homologous. In infection experiments, DNA A1 efficiently transreplicated DNA B3 but failed to replicate DNA B1 and DNA B2. This suggested that further DNAs A were present (Jovel *et al.*, 2004). Here, we describe a series of experiments combining PCR, RFLP, cloning and sequencing techniques that allowed the identification of two new DNAs A (DNA A2 and DNA A3) of SimMV, which remained undetectable in the *S. micrantha* plants for more than twenty years. The series of experiments described here have the potential to be used as diagnostic tools for the study of mix infections originating diseases in crop fields.

## Material and Methods

### Plants and viruses

*S. micrantha* plants harbouring the Sida micrantha mosaic viruses have been described (Abouzid and Jeske, 1986) and will be referred to in this paper as “original *S. micrantha* plants”. Some viruses forming the SimMV complex have also been described (Jovel *et al.*, 2004; Jovel *et al.*, 2007).

### Extraction of Nucleic Acids

One young and symptomatic leaf was shock frozen in liquid nitrogen, ground to powder, and shaken for 15 min at room temperature (RT) with 500 μl homogenization buffer (100 mM Tris– HCl, pH 7.5; 1 mM EDTA; 100 mM NaCl; 0.6% SDS; 100 mM DTT) plus an equal volume of phenol-chloroform (PC, 9:1). The suspension was centrifuged for 5 min at RT (14000 rpm). The supernatant was extracted with a volume of chloroform for 5 min, centrifuged 5 min as above, and finally precipitated with one tenth of 3M NaAc (pH 4.8) and two volumes of ethanol (EtOH), overnight at – 20°C. After centrifugation (30 min, 14000 rpm, 4°C), the pellet was washed with 0.5 ml 70% EtOH, recovered by centrifugation (5 min, 14000 rpm, RT), dried at RT and re-suspended in 50 μl of 1x TE buffer (10 mM Tris-HCl, 1 mM Na-EDTA, pH 7.5).

### Biolistic inoculation

Total nucleic acids extracted from the original *S. micrantha* plants were bombarded onto six-leaf-stage *M. parviflora* or *Nicotiana benthamiana* plants as described by Zhang *et al.* (2001).

### Polymerase chain reaction (PCR)

PCR was performed in the presence of 10 pmol of each primer, 20 mM of each dNTP, 1.5 mM MgCl_2_, 10 mM Tris-HCl (pH 7.5), 0.25 units of Taq DNA polymerase (Qiagen, Germany) and 100 ng of total DNA as template, in a final volume of 50 μl. The following cycling program was used: [5’, 95°C; 1’, 55°C; 1.5’, 72°C; 1X / 1’, 95°C; 1’, 55°C; 1.5’, 72°C; 30X / 1’, 95°C; 1’, 55°C; 10’, 72^?^C; 1X].

### Southern blot and hybridization of nucleic acids

Total nucleic acids from infected plants were loaded onto a 1% agarose gel containing 0.4 μg/ml EtBr and 0.5x TBE buffer (8.9 mM Tris-HCl, 8.9 mM boric acid, 2 mM EDTA) and electrophoresed in the presence of 0.5x TBE. Afterwards, the gel was shaken for 20 min in alkaline transfer solution (0.6 M NaCl, 0.4 M NaOH) and the DNA transferred to a nylon membrane (Hybond +) by Southern blotting in the presence of the same solution.

### Labelling of probes and hybridization

The AC1/AC3 (primers SmRep-V and SmRep-C; table 1) and BC1 (primers B3C1-V and B3C1-C; Jovel et al., 2004) regions, for DNA A and DNA B, respectively, were PCR-amplified and precipitated. For probe labelling, eight μg of DNA were denatured (95°C, 10 min), quickly chilled, and incubated with 4 μl of DIG-Prime kit (Roche, Mannheim, Germany), overnight at 37°C. In order to remove non-incorporated nucleotides, the labelled DNA was precipitated for 2h in the presence of 40 mM LiCl, 10 μg of yeast tRNA, and 2 volumes of EtOH. The membrane was prehybridized for 3h at 42°C in pre-soak solution [1% glycine, 5x SSPE (20x SSPE: 3M NaCl, 0.2 M NaH_2_PO^-4^H_2_0, 0.02 M Na-EDTA, pH 7.4), 40μg/ml fish sperm, 100x Denhardt (0.5% BSA, 0.5% Ficoll 400, and 0.5% PVP 4000), 50% deionised formamide, 0.12% SDS], and then hybridized overnight in hybridization solution (in which the glycine of the pre-soak solution was replaced by 10% dextran-sulphate and the labelled probe). Both, pre-soak and hybridization solutions were heated at 80°C for 10 min prior to use.

**Table 1.**
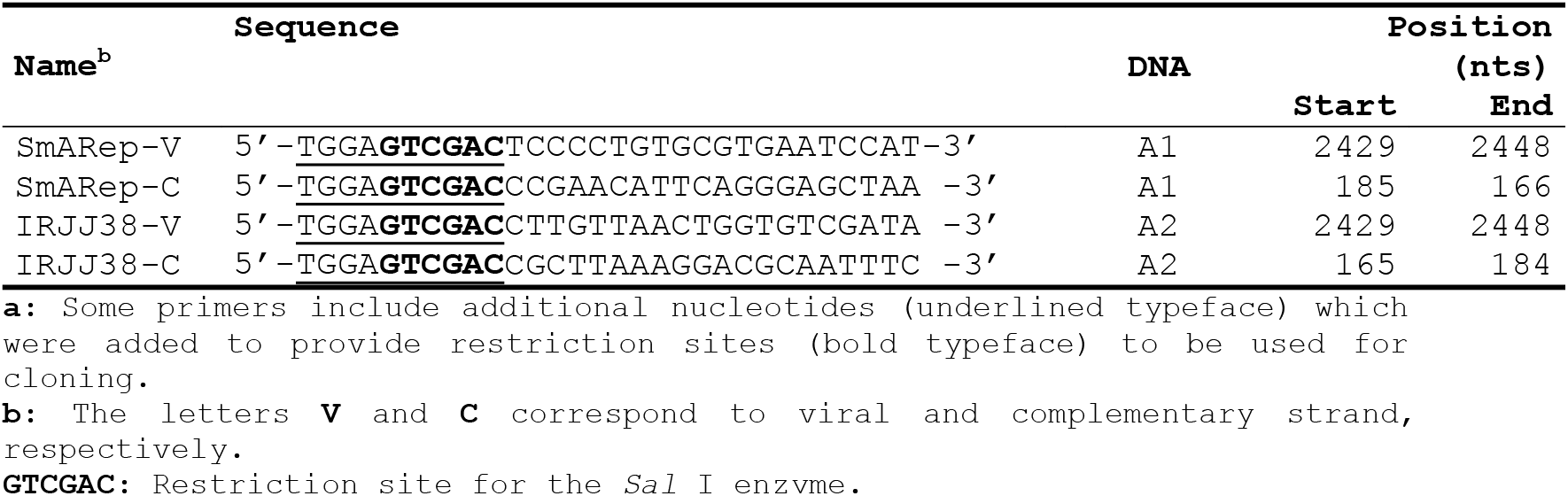
Primers used to amplify the sequences used as probes.

The membranes were washed for three 15-min periods at 42°C in the presence of 2x SSPE and 0.3% SDS and then soaked for 3 min in buffer 1 (0.1 M maleic acid, 0.15 M NaCl, pH 7.5). Blocking of membranes was carried out in buffer 2 (buffer 1 plus 1% blocking reagent (Roche)) for 30 min. Digoxigenin detection was for 30 min in a 1:10, 000 dilution of anti-digoxigenin AP (DIG Luminiscent detection kit, Roche) in buffer 2. Three posthybridization washes were performed in buffer 1 plus 0.03% Tween-20 for 15 min. Finally, membranes were incubated for 5 min in buffer 3 (100 mM Tris-HCl, pH 9.5; 100 mM NaCl; 50 mM MgCl_2_), and finally for 5 min in a 1:100 dilution of the substrate CSPD or CDP Star (Roche) in buffer 3.

### Restriction fragments length polymorphisms (RFLP)

Polyacrylamide gels contained 8% acrylamid/bisacrylamide (30:0.8; Roth, Karlsruhe, Germany), 1x TBE, 0.14% ammoniumperoxodisulphate (APS), and 0.014% tetramethylethylendiamin (Temed). 500 ng of the PCR product were digested with 5 units of *Alu* I for 3 h at 37°C and resolved by polyacrylamide gel electrophoresis (PAGE) in the presence of 1x TBE buffer at 60 V.

## Results

DNA from six *S. micrantha* plant lines originally imported from Brazil to Germany in 1977 was used to clone, stepwise over several years, DNA A1, DNA B1, DNA B2 and DNA B3 of SimMV (Jovel *et al.*, 2004). On detached leaves and plants of *N. benthamiana,* DNA A1 was able to trans-replicate only its cognate DNA B3, but not DNA B1 or DNA B2. This led us to hypothesize that either additional DNAs A had to be present in the viral complex or that DNA B1 and DNA B2 could have vanished over the years of vegetative propagation of *S. micrantha* plants in the apparent absence of cognate DNAs A. In order to scrutinize for the presence of the distinct DNAs B cloned over the time, the BC1 open reading frame (ORF) of DNA B1, B2 and B3 were searched with specific primers (Jovel *et al.*, 2004). The sequence of the primers for DNA B1 and DNA B2 is identical. The obtained PCR products were digested with *Alu* I and the resulting polymorphisms analyzed by PAGE. Five out of six plant lines analyzed contained both DNA B2 and DNA B3 (Fig. 1A, B) but not DNA B1 (data not shown). In plant line I-14, the primers for DNA B1/B2 amplified a fragment that resulted in an *Alu* I digestion pattern that did not correspond neither to DNA B1 nor to DNA B2. Primers for DNA B3 amplified only a very faint band. Further analyses to discern the sequence of such amplicon were not done.

**Figure 1.**
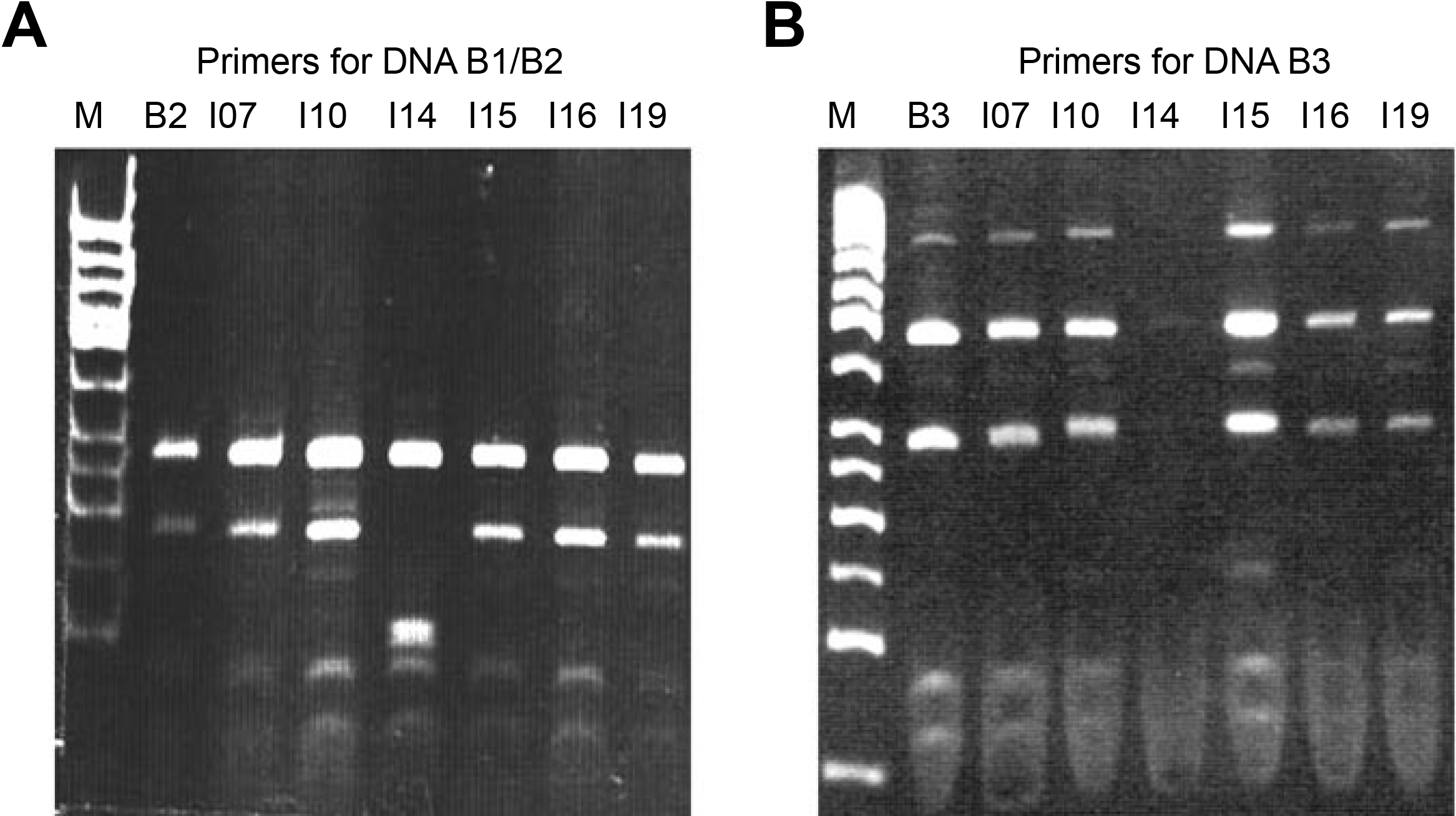
PCR/RFLP analysis of the BC1 ORF of DNA B2 (**A**) and DNA B3 (**B**) in the original *S. micrantha* plant lines I–07, I–10, I– 14, I–15, I–16, and I–19. Five hundred ng of PCR product were digested with 0.5 Units of *Alu* I for 3 h at 37°C, resolved by PAGE, stained with EtBr and visualized under UV light. The reference pattern of DNA B2 (A) and DNA B3 (B) were included.

Numerous trials in search for a further DNA A in the original *S. micrantha* plants were unsuccessful. (**1**) Supercoiled DNA purified from these plants was digested with frequently-cutting enzymes, cloned in *E. coli* and sequenced but the sequence of all clones corresponded to those obtained previously (DNA A1, DNA B1, DNA B2 and DNA B3). (**2**) Amplification of the IR of DNAs A, by using primers SmAIR-V and SmAIR-C (Jovel *et al.*, 2004) located at highly conserved motifs of the AC1 and AV1 ORFs, and digestion of the resulting PCR products with different restriction enzymes did not reveal additional polymorphisms. An unintended transmission of begomoviruses from Sida plants from our collection to experimental *M. parviflora* plants provided the opportunity to find new geminiviruses previously unidentified as well as pseudo-recombinants. This is described hereafter.

Since geminiviruses are transmitted by the whitefly *Bemisia tabaci,* which is not present in Germany, experimental *M. parviflora* plants that had been bombarded with clones of SimMV DNA A1 in combination with DNA B1, DNA B2 or DNA B3, were placed in the same cabin with several geminivirus-infected *Sida* plants collected in different countries of Latin America and Africa. Two weeks post inoculation, symptomatic plants were observed in the three groups of inoculated plants. High levels of viral DNA in systemic leaves were unexpected (Fig. 2A), especially in case of plants bombarded with DNA A1/B1 or DNA A1/B2, since in transient assays DNA A1 was unable to trans-replicate DNA B1 or DNA B2 to detectable levels.

**Figure 2.**
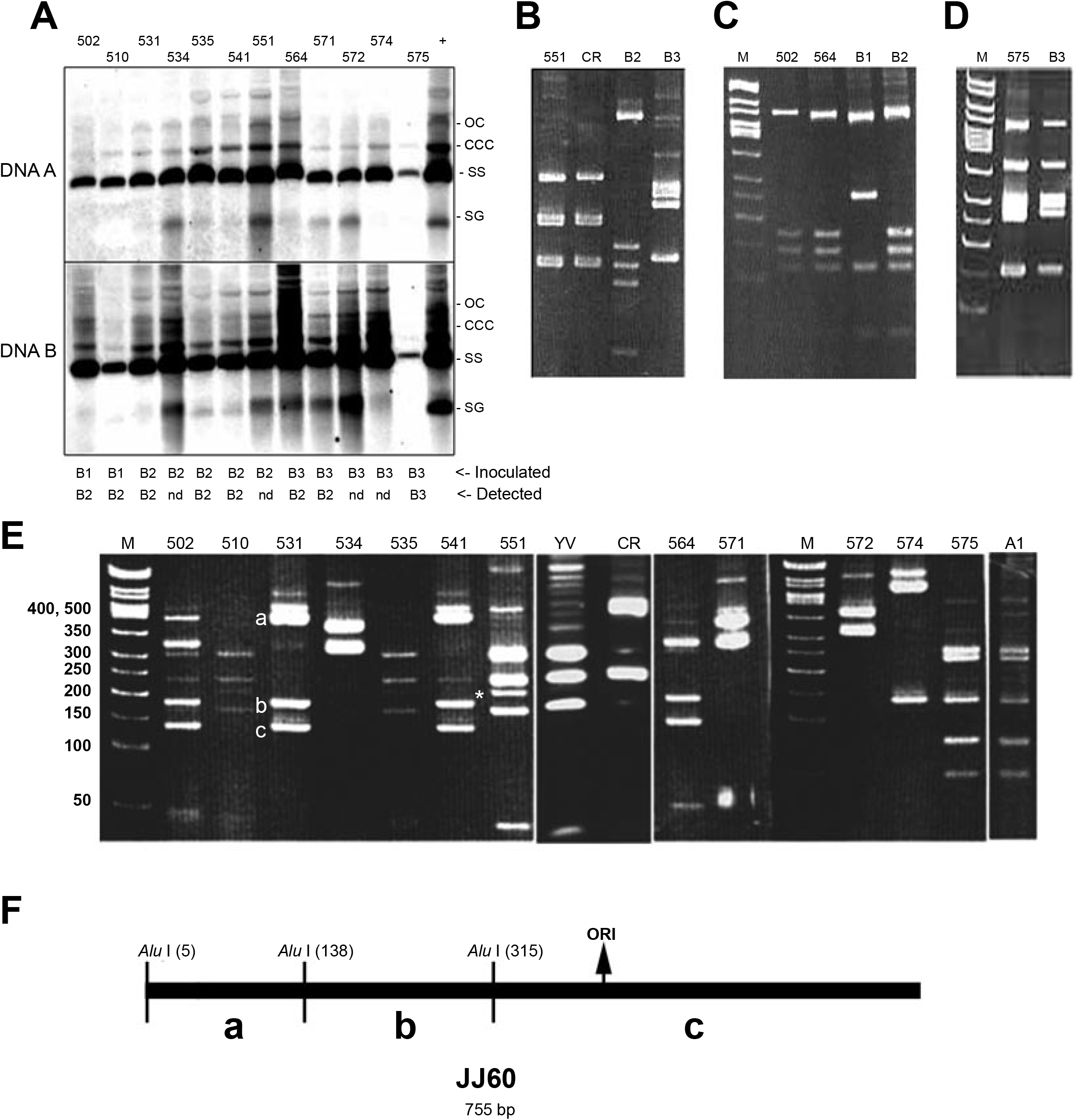
(**A**) Detection of viral DNA in *M. parviflora* plants by Southern blotting and hybridization. One μg of total nucleic acids was loaded for each of the twelve symptomatic plants (502 … 575) and hybridization was carried out using a DNA A-specific probe containing a fragment spanning the AC1/AC3 ORFs or a DNA B-specific probe containing the BC1 ORF. (+): one μg of total nucleic acids from original *S. micrantha* plants. Viral DNA forms are indicated at the right border (oc: open circular, ccc: covalent closed circular, ss: single stranded, sg: subgenomic). Both the DNA B that was inoculated in the experimental contaminated plants and that detected by the *Alu* I-RFLP analysis (data not shown) are listed at the bottom of the picture. (n.d.: not determined). (**B**) The IR of DNA B was amplified from plant 551 (primers SmB1/2IR-V and SmB1/2IR-C) and compared to the corresponding region in SiGMCRV DNA B (CR), and SimMV DNAs B2 and B3. (**C**) The IR of plants 502 and 564 (amplified as in B) was compared to the corresponding region in DNA B1 (B1) and DNA B2 (B2). (**D**) The IR of plant 575 (primers SmB3IR-V and SmB3IR-C) was compared to the corresponding region in DNA B3. No other primer combination produced any amplicon in B, C and D. (**E**) Detection of polymorphisms in the IR of DNAs A. Fragments were amplified by using primers SmAIR-V and SmAIR-C. The digestion pattern of DNA A1 of SimMV (A1) as well as those of SiYVV (YV) and SiGMCRV (CR) were included as controls. (**E**) The restriction map of the IR of clone JJ60 is included and the corresponding fragments indicated by letters.

Therefore, to scrutinize the source of infection, the intergenic region of DNA B1, B2 and B3 was amplified with suitable primers (Jovel *et al.*, 2004). An *Alu* I-RFLP analysis was performed and the products compared with reference plasmids. Surprisingly, the primers for DNA B1/DNA B2 were able to amplify the IR of the DNA B of *Sida golden mosaic Costa Rica virus* (SiGMCRV) in one of the infected plants (plant 551; Fig. 2B). DNA B2 of SimMV was present in several plants (502, 510, 531, 535, 541, 564, and 571; data not shown), including some in which DNA B2 had not been inoculated, for instance plants 502 and 564 (Fig. 2C). The primers for DNA B3 amplified a fragment of the expected length only in plant 575, which was verified as DNA B3 by RFLP (Fig. 2D). This plant had certainly been inoculated with DNA A1/B3. None of the plants contained DNA B1 (data not shown). In summary, the obtained RFLP results were not compatible with the inoculation scheme (Fig. 2A, bottom), which led us to search for possible sources of contamination. Closer inspection of the plants revealed that *B. tabaci* had been introduced with unknown origin in our greenhouse. Therefore, we concluded that the transmission of SimMV DNAs A was probably accomplished by whiteflies and not by contaminated bombardment devices. This obvious accident, however, buried an opportunity to identify new DNAs A, as described bellow.

If DNA B2 was present in some of these plants (Fig. 2C), the DNA A responsible for the trans-replication of DNA B2 should also have been transmitted from original *S. micrantha* plants to *M. parviflora.* Therefore, we proceeded to analyze the polymorphisms of the IRs of the DNAs A in the *M. parviflora* plants. The *Alu* I-RFLP analysis revealed a high diversity of DNAs A that had been transmitted by *B. tabaci* onto such plants (Fig. 2E), but surprisingly only one of them contained the DNA A1 of SimMV, the plant 575, which also contained DNA B3 (Fig. 2D). This plant contained only low concentration of viral DNA (Fig. 2A), exhibited mild symptoms, and recovered after six weeks.

A peculiar combination was found in plant 551. As mentioned above, the DNA B of SiGMCRV was clearly identified in this plant (Fig. 2B). However, after the *Alu* I-RFLP analysis (Fig. 2E), cloning and sequencing of twenty IRs clones from this plant, the DNA A was identified as *Sida yellow vein virus* (SiYVV), from Honduras (Frischmuth *et al.*, 1997) (compare in Fig. 2E, sample 551 with YV, which corresponds to the IR from a plasmid containing the DNA A of SiYVV). An atypical band (* in Fig. 2E) which does not correspond to the restriction pattern of SiYVV nor to that of SiGMCRV (compare sample 551 with lane CR, which corresponds to the IR from a plasmid containing the DNA A of SiGMCRV) was detected. We assumed it represents an unspecific PCR product. The symptoms in this plant were severe and included excessive proliferation of leaves, mottling, and stunting. Indeed, the plant’s height never reached more than 7–8 cm, compared to 50 cm on average in control plants (Fig. 3B). Excessive proliferation of leaves had also been observed in several species of plants experimentally infected with DNA A and DNA B of SiGMCRV (Jovel and Ruiz, 2004), suggesting that SiGMCRV DNA B is responsible for the induction of this symptom. However, extreme stunting is neither typical for SiGMCRV nor for SiYVV, indicating that the pseudo-recombinant virus possessed increased pathogenicity.

**Figure 3.**
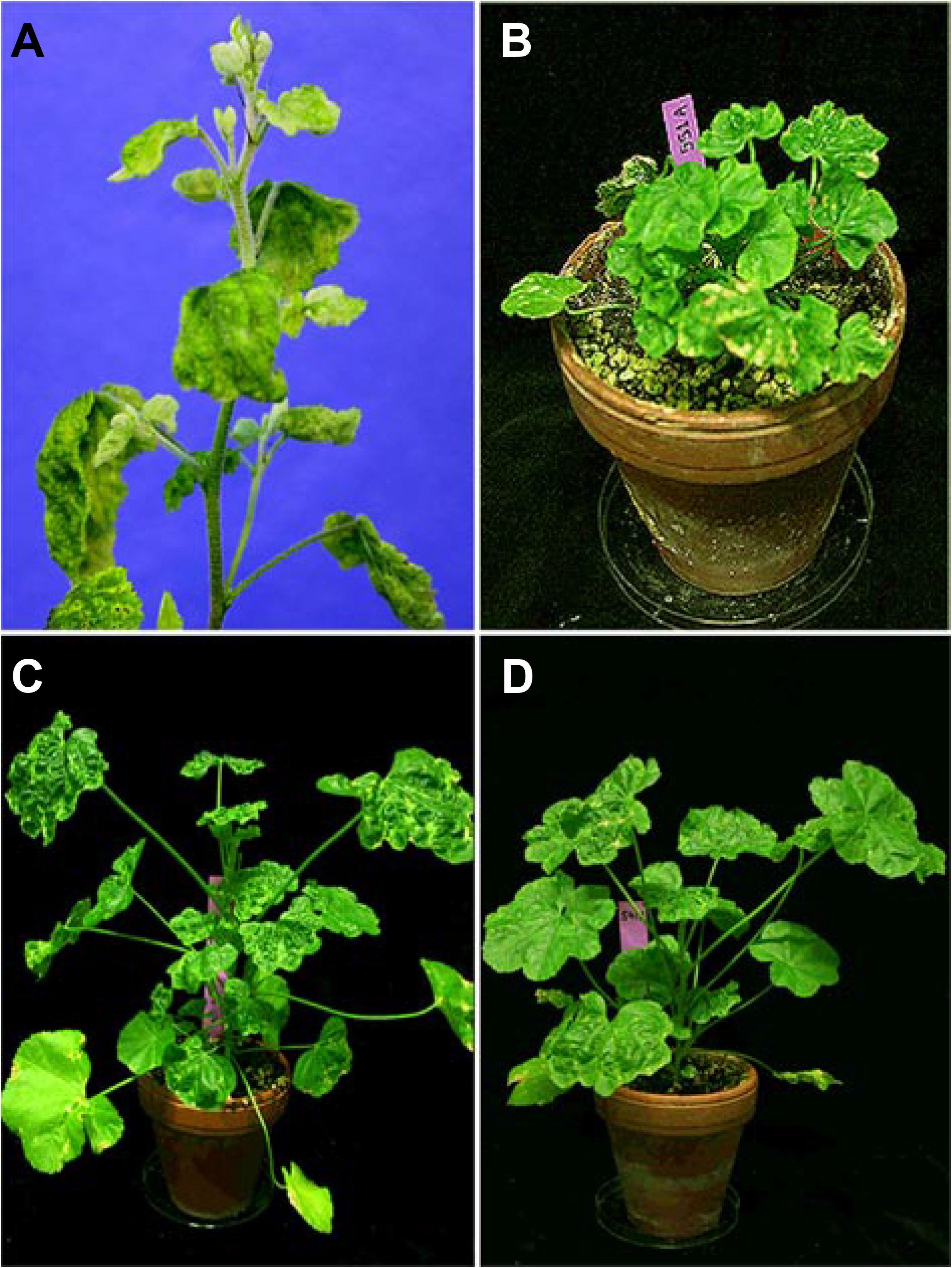
(**A**) Symptoms induced by the SimMV complex in the original *S. micrantha* plants. (**B**) Plant 551 containing the pseudorecombinant formed by SiYVV DNA A and SiGMCRV DNA B. (**C**) Plant 564 in which DNA A2 and DNA B2 of SimMV were detected. (**D**) Plant 541 containing the psedorecombinant formed by the putative DNA A from Macropthilium mosaic virus and SimMV DNA B2. Symptoms in **C** and **D** were very similar in strength. It has been shown that DNA B is responsible for symptom induction (Pascal *et al*., 1993).

Plants 502 and 564, harbouring DNA B2 (Fig. 2C), did not contain DNA A1, but new DNAs A (Fig. 2E). Plant 502 contained an additional DNA A, apparently corresponding to SiYV, which was also present in plants 510 and 535. Plant 564 apparently contained one single DNA A-like molecule and for this reason was selected for further experiments. Symptoms in plant 564 were severe and included wrinkling of leaves and mosaics (Fig. 3C). Supercoiled DNA from this contaminated *M. parviflora* plant was extracted and bombarded onto *N. benthamiana.* From these latter, the IR of a DNA A was amplified by PCR, cloned in *E. coli* (clone JJ38) and sequenced. The CR of this DNA A-like molecule was identical to that of DNA B2. The full-length of this new DNA A (DNA A2) was cloned and sequenced as described in Jovel *et al.* (2004). Plants 531 and 541 (Fig. 2E) exhibited an *Alu* I– restriction pattern similar to that of DNA A2 (plant 564), and also contained DNA B2 (data not shown). To elucidate if a third DNA A was involved in the trans-replication of DNA B2 in these plants, or whether a mutant form of DNA A2 was the origin of such *Alu* I-digestion pattern, a large number of clones comprising this IR were generated and sequenced. The sequences of all clones were identical, and 91% homologous to that reported for Macropthilium golden mosaic geminivirus (accession number AF098940), originally isolated from *Macropthilium lathyroides* in Jamaica. The restriction pattern of one representative clone sequenced is shown in Fig. 2F to demonstrate the congruency of the restriction pattern with the sequence. This suggested that a natural pseudo-recombination event took place between DNA B2 and this additional DNA A. The symptoms induced by this pseudo-recombinant were very similar to those found in plant 564, which was infected with DNA A2/B2. This is consistent with the notion that in begomoviruses the DNA B genomic component is responsible for the induction of symptoms (Pascal *et al.*, 1993). It is very likely that this newly– identified molecule is the parental form of DNA A2, or vice versa.

Once the full length of DNA A2 was obtained, trans-replication experiments combining SimMV DNA A1 and DNA A2 with the three DNAs B (B1, B2 and B3) were conducted. DNAs A1 and A2 were able to trans-replicate specifically DNA B3 and DNA B2, respectively (Fig. 4). Trans-replication experiments were conducted on detached leaves and whole plants of *N. benthamiana.* In detached leaves, DNA A1 was able to trans-replicate DNA B3 and no traces of single-stranded or covalently-closed circular viral DNA forms were detected for DNA B1 and DNA B2 (Fig. 4A, left panel). Analogously, DNA A2 specifically trans-replicated DNA B2 (Fig. 4A, right panel). In both cases, relatively high amounts of single-stranded DNA were found, but other forms were scanty. However, when the cloned DNAs were bombarded on *N. benthamiana* plants, DNA A1/B3 and DNA A2/B2 accumulated high amount of all replicative forms (Fig. 4B). In both cases, symptoms induced in *N. benthamiana* were mild: plants were slightly stunted and showed a yellowish mosaic. In some plants infected with both DNAs A1/B3 and DNAs A2/B2, symptoms were apparently more severe than in single infections (Fig. 4D). Interestingly, DNA A1/B3 accumulated to higher levels in *N. benthamiana.* It is possible that these begomoviruses cooperatively interact to induce severe symptoms; however, a more meticulous analysis is still required. To verify that the inoculated DNAs were responsible for the induction of symptoms and detected viral DNA, the IR of DNA A1, DNA B3, DNA A2 and DNA B2 were amplified and digested with *Alu* I. When compared to reference plasmids, we found that effectively the inoculated DNAs were present in the infected plants (Fig. 4C).

**Figure 4.**
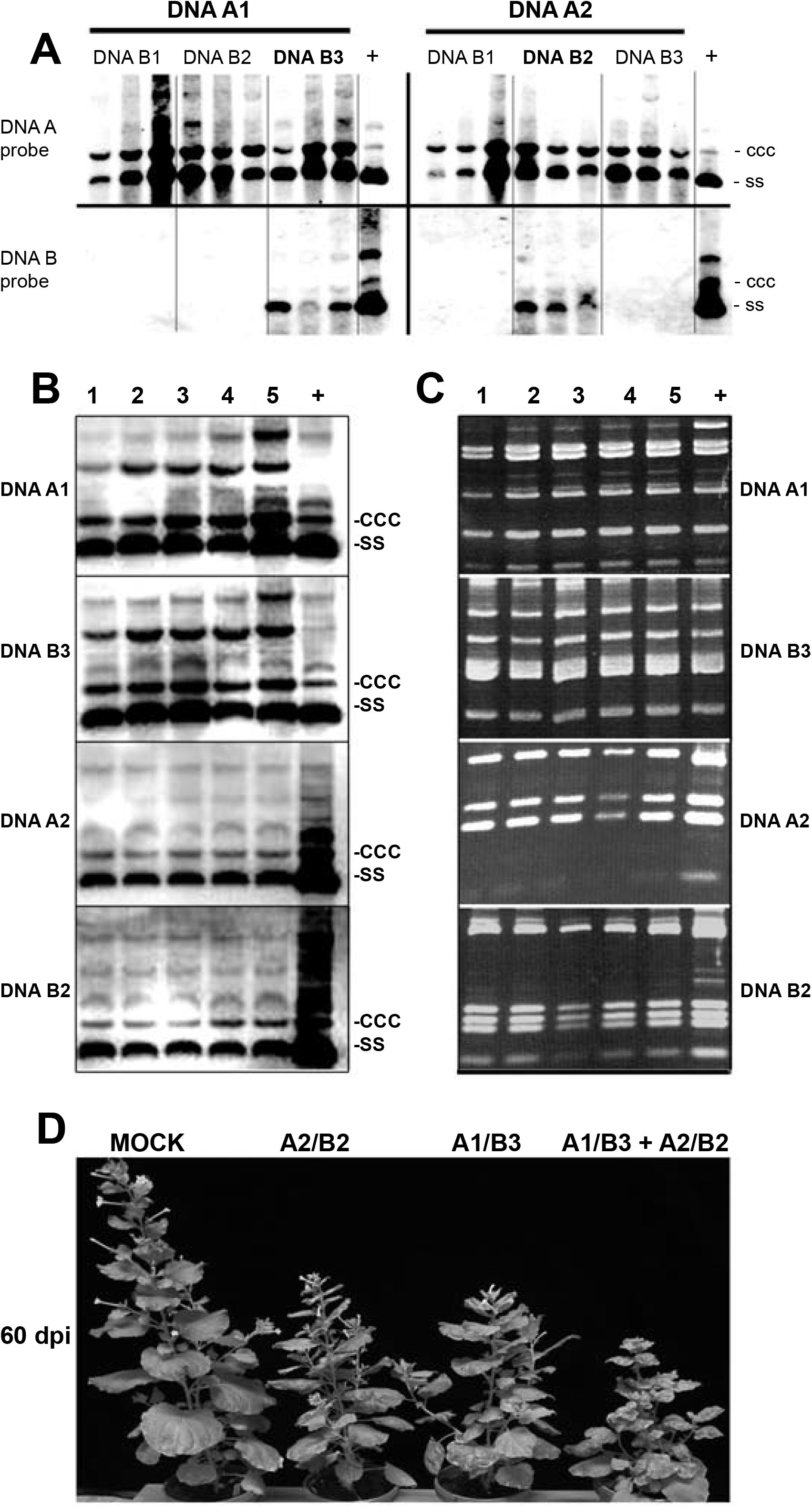
(**A**) Analysis of the ability of DNA1 and DNA A2 to transreplicate each of the DNAs B in transient experiments. DNA A1 (left panel) and DNA A2 (right panel) were co-bombarded onto detached *N. benthamiana* leaves either with DNA B1, DNA B2 or DNA B3, as indicated. DNA of original *S. micrantha* plants was used as positive control (+). A DNA A1 and a B3C1 probe, which in a preliminary experiment were able to hybridize non– specifically against every DNA A and DNA B, respectively, were used under conditions of low stringency as described in Sambrook and Russell (2001). (**B**) One μg of DNA from five DNA A1/DNA B3-and five DNA A2/DNA B2-infected *N. benthamiana* plants was hybridized against the probes described above, under low stringency conditions. One μg of DNA from original *S. micrantha* plants was used as positive control (+). (**C**) DNA from the same plants in panel B was used to amplify the IR of each DNA. The PCR products were digested with *Alu* I. As a control (+) the IR of each DNA was amplified from the corresponding plasmids and digested as above. (**D**) Symptoms induced in *N. benthamiana* by DNAs A1/B3, DNAs A2/B2 or a combination of both. Plants in D were inoculated by biolistic bombardment of cloned DNAs. Photo was taked 60 days post inoculation.

One critical question raised by the discovery of DNA A2 was whether it had really been transmitted from the original *S. micrantha* plants, where it could not be detected before, or was created by a recent recombinational or mutational event imposed by the new cellular environment in *M. parviflora.* To investigate this question, we identified a small fragment in the IR of DNA A2 that was dissimilar enough from DNA A1 to be used as selective primer binding site (IRJJ38-V and IRJJ38-C; table 1). PCR reactions were performed as described but with increased annealing temperature of 62°C. Under such conditions, these primers selectively amplified the desired fragment from the plasmid containing the IRs of DNA A2 but not that of DNA A1 (Fig. 5A). By using such specific primers, we were able to amplify the corresponding fragment from all *S. micrantha* plants and the *Alu* I-restriction pattern showed that is was identical to that of DNA A2 (Fig. 3B, C).

**Figure 5.**
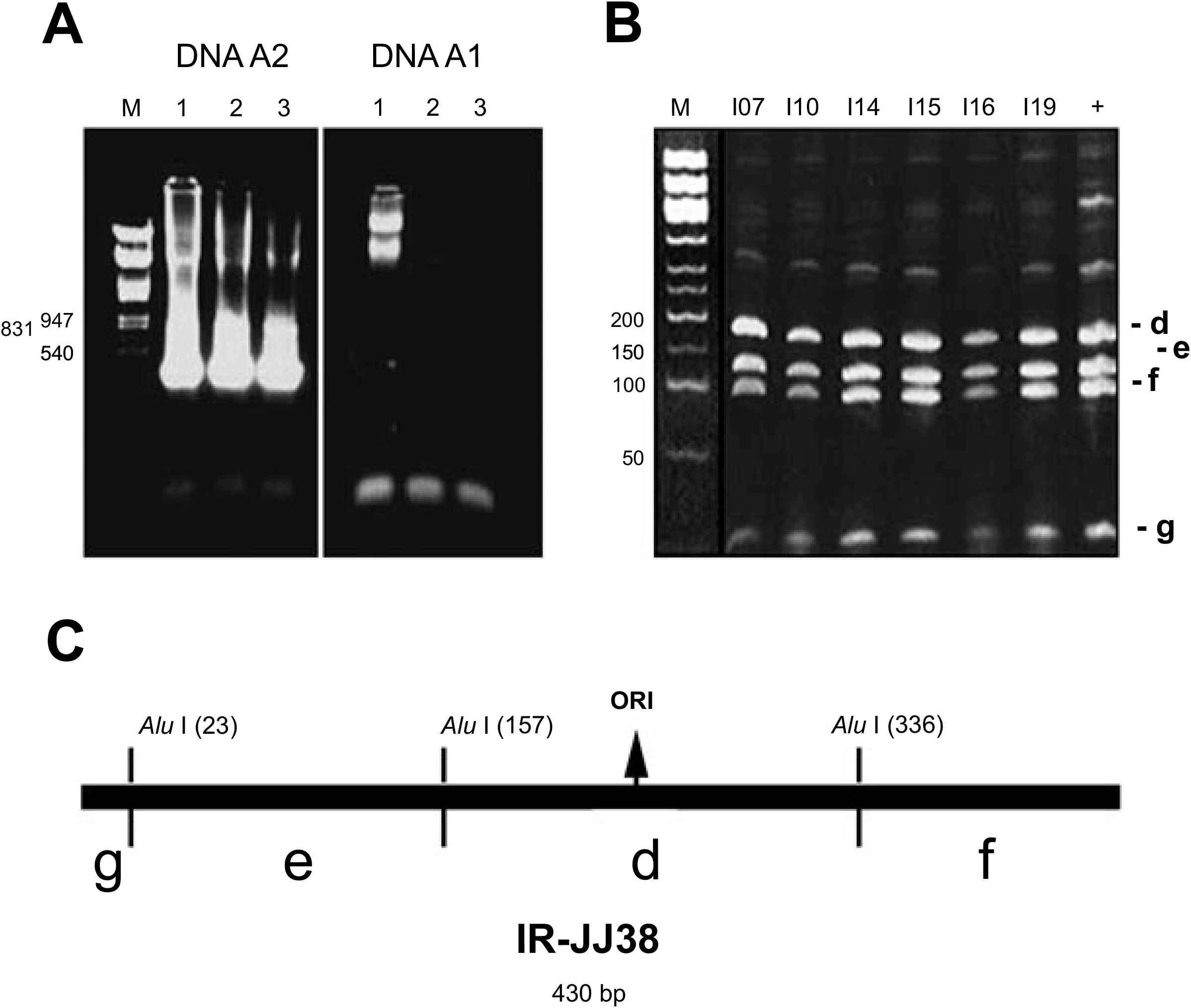
Identification of DNA A2 in the original *S. micrantha* plants. (**A**) A fragment within the IR of DNA A2 was specifically amplified by using primers IRJJ38-V and IRJJ38-C (Table 1), which were able to discriminate between the plasmids containing DNA A1 and DNA A2. (**B**) The *Alu* I restriction pattern was identical to that of DNA A2, and was found in all six original *S. micrantha* plant lines. (**C**) The restriction map of the amplified fragment is included for comparison. Ori: origin of viral replication.

Why DNAs A2/B2 were transmitted to a larger number of *M. parviflora* plants than DNAs A1/B3? Does *B. tabaci* transmit DNAs A2/B2 preferentially over DNAs A1/B3? Was *M. parviflora* a differential host for both viruses? To answer these questions, total DNA from the original *S. micrantha* plants was extracted and bombarded onto five to six-leaf-stage *M. parviflora* plants. Two weeks after bombardment, seven out of eight plants became infected and exhibited symptoms visually indistinguishable from those in the *M. parviflora* plants accidentally contaminated with DNAs A2/B2 (see Fig. 3C). Total DNA from these plants was extracted, digested with *Hae* II, blotted, and hybridized against a DNA A2 probe without CR, thereby preventing cross-hybridization with DNA B2. DNA A2 was found as the major component of these infected plants, suggesting that DNAs A2/B2 are better adapted to this host plant than DNAs A1/B3. Another possibility is that transmission of DNAs A2/B2 by *B. tabaci* may be more efficient. It has been demonstrated that, in bipartite geminiviruses, the coat protein is involved in transmission by whiteflies, and specific amino acids that determine transmissibility have been delineated (Hohnle *et al.*, 2001). Amino acid sequence analysis of the coat protein of DNA A1 and DNA A2 showed that none of these molecules contain mutations that could affect transmissibility by *Bemisia tabaci.*

Finally, as described in Jovel *et al.*, (2004), the diversity of DNAs A in the original *S. micrantha* plants brought from Brazil to Germany more than two decades ago, was explored by PCR/RFLP analysis. The results showed that from the seven plant lines analyzed, only lines I-14 and I-16 exhibited additional restriction bands, which suggested the presence of further DNAs A (Jovel *et al.*, 2004). The IR of DNAs A in such plant lines was PCR amplified (primers SmAIR-V and SmAIR-C; Jovel *et al.*, 2004), cloned and sequenced. No novel sequences were detected in plant line I–16 and all clones corresponded to DNA A2.

However, I-14 contained an unknown DNA A-like molecule (clone JJ95) apparently with a recombinant IR (Fig. 7A). The region upstream of the TATA box for complementary strand transcription, including a ~250-nt-long stretch of the AC1 protein shared 95% homology with *Tomato rugose mosaic virus* (ToRMV-[Ube]), whereas the ~330 nts downstream, including ~15 nts of the coat protein gene, were 96% homologous to DNA A2 of SimMV. Again, as in the case of DNA B2, a very short motif (TATATA, Fig. 7A) seems to have served as the point for matrix exchange during replication. As a consequence of the recombination, this new molecule possesses and AC1 binding site (**GGTAG**ttat**GGTAG**) different from that of DNA A2, but surprisingly identical to that of DNA B1. Sequence comparisons identified a ~140-nt-long CR between DNA A3 and DNA B1 (Fig. 7B), suggesting that they are cognate components of a third begomovirus in the SimMV complex. We have called this new molecule DNA A3.

## Discussion

### Mixed infections originating a syndrome

Cloning and characterization of the SimMV has been a 25–years–long task (Jovel *et al.*, 2004). During that period, several viruses infecting various species of *Sida* from Costa Rica (Höfer *et al.*, 1997b) and Honduras (Frischmuth *et al.*, 1997) were cloned and sequenced, evidencing that the heart of the problem was not the plant type (Malvaceae) but rather the virus itself or its interaction with the host plant.

Several factors boost the emergence of viral complexes in the *Begomovirus* genus. Firstly, the ample host range of their insect vector *(B. tabaci)* implies that begomoviruses are transmitted to different plant species with variable cellular environments where they concur with other geminiviruses. When this happens, molecular recombination, pseudo-recombination and mutation may create novel viral forms better adapted to the new cellular conditions, which do not necessarily imply that all parental forms vanish. So might emerge a viral complex.

The Sida micrantha mosaic disease was found to be associated with a complex of functionally independent begomoviruses (Jovel *et al.*, 2004). Thus, the problem was to elucidate how many viral entities were associated to determine such syndrome and, more critically, which begomovirus exerts the predominant effect to produce the disease. The aetiology of the *Sida micrantha* mosaic still remains a mystery. Although three independent begomoviruses have been isolated, the retro-introduction of them into *S. micrantha* plants did not succeed so far. We do not rule out that additional viruses could be present in the Sida mosaic plants brought from Brazil. It is indeed very likely that a hidden variability might be underestimated since geminivirologist are used to concentrate our attention on bands of DNA in the separating gels and neglect the less prominent but, possible, the majority of the molecules in the Gaussian distribution. Under the concept of quasispecies of viruses (Eigen, 1993), however, this circumstance should be considered to define the causative agents of syndromes. Perhaps it is therefore necessary in future to combine many independent molecular clones of all DNA components to restore the biological variability and to reproduce the full syndrome in Sida plants.

Plant cloning and especially molecular cloning are always selection processes resulting in the singular representation of viruses or DNA molecules of a complex mixture. In the best case, they reflect the most frequent component which is not necessarily the most effective one. In the worse case, if the clone is harmful to or unstable in *E. coli,* molecular cloning might select for clonability rather for than representative members of the original pool. Reviewing the twenty-five years of solving the *S. micrantha* mosaic conundrum, the latter explanation seems to hold true for this begomoviral complex. This may be particularly valid for DNA A2, which has been clonedas the last molecule only after biological purification by passaging through *M. parviflora* and *N. benthamiana* plants, although it appeared as the major viral component in *S. micrantha*.

The cloned versions of DNAs A1/B3 and DNAs A2/B2 exhibited some interesting biological differences. For example, DNAs A2/B2 were found in 7 out of 12 accidentally contaminated *M. parviflora* plants (compared to one single plant infected with DNAs A1/B3) suggesting that the former virus is better adapted to *M. parviflora* than DNAs A1/B3. Alternatively, *B. tabaci* might preferentially acquire and/or transmits DNAs A2/B2. We reject the latter explanation since inoculation of total DNA from *S. micrantha* plants onto *M. parviflora* resulted also in a higher accumulation of DNA A2 compared to DNA A1 (Fig. 6). Moreover, previous studies have established that the different biotypes of *B. tabaci* show no preferences for the transmission of begomoviruses (Bedford *et al.*, 1994).

**Figure 6.**
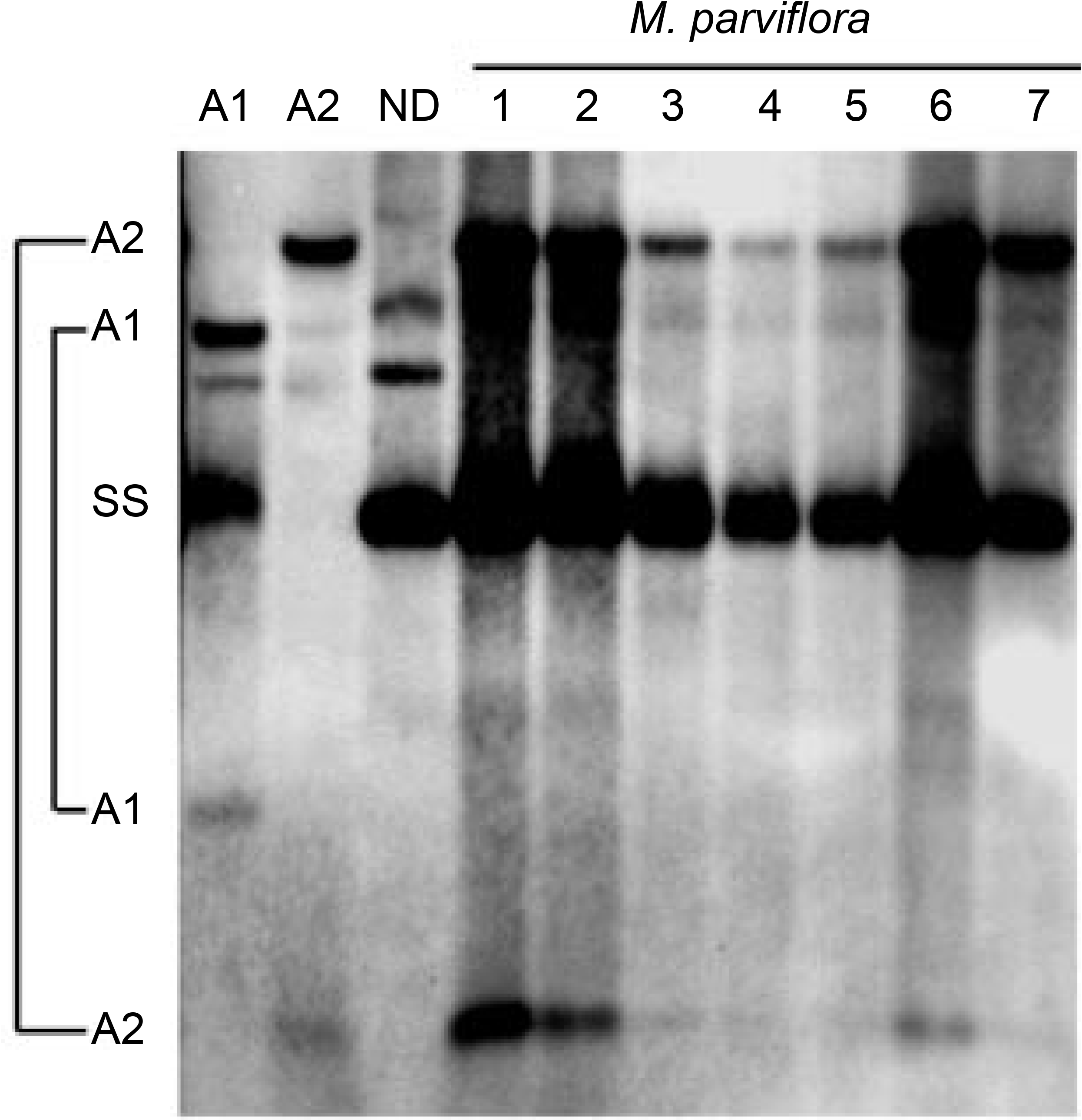
Southern blot analysis of DNA from seven *M. parviflora* plants infected with total nucleic acids from the original *S. micrantha* plants. DNA was digested with *Hae* II and hybridized with a DNA A2 probe without CR. A1: DNA from a *N. benthamiana* plant infected with DNAs A1/B3. A2: DNA from a *N. benthamiana* plant infected with DNAs A2/B2. ND: Non-digested DNA from *S. micrantha* plants was included as controls to identify nondigested forms of the viral DNA.

**Figure 7.**
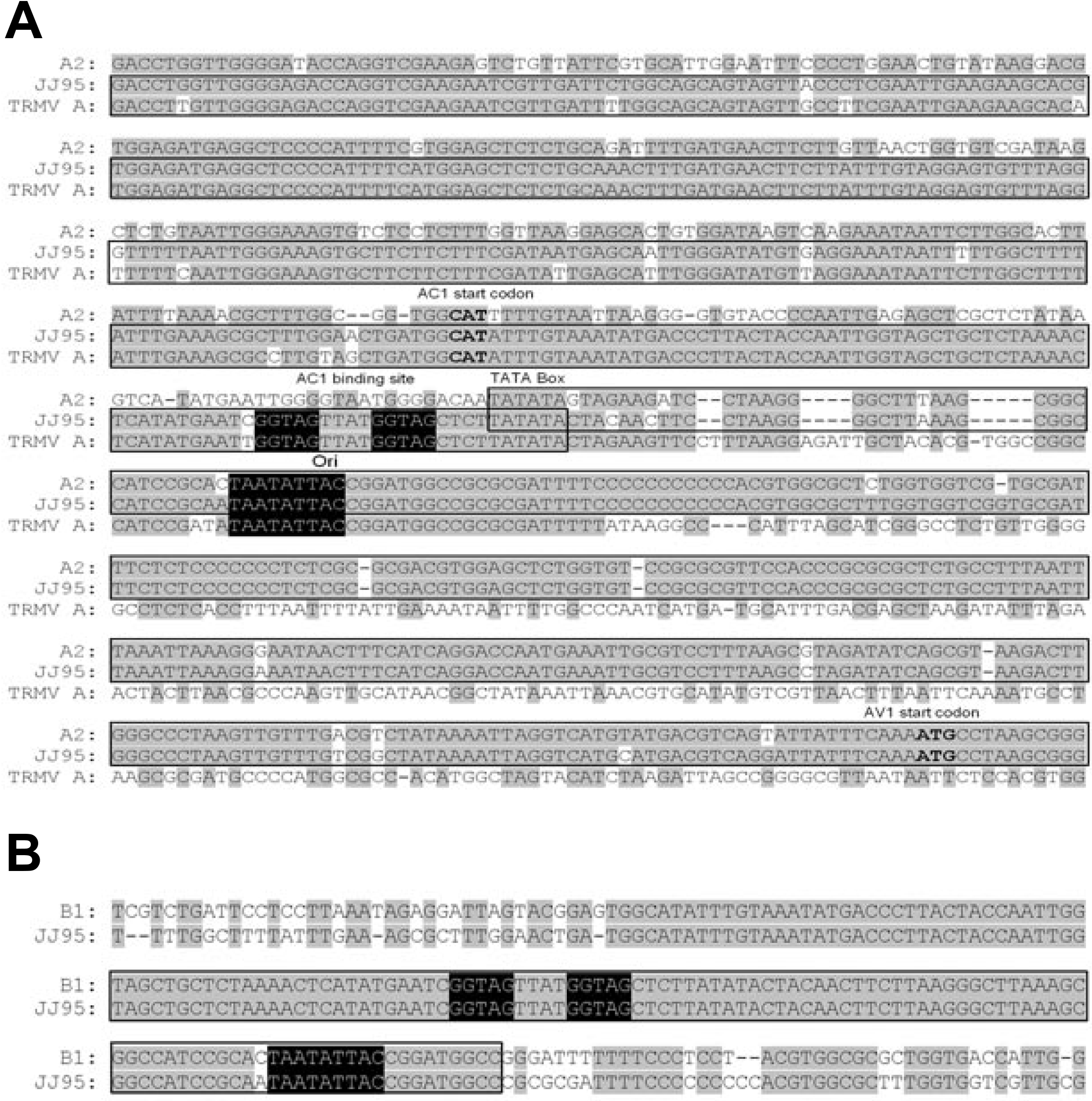
(**A**) Alignment of the IRs of SimMV DNA A2, ToRMV-[Ube] (accession number AF291705) DNA A and clone JJ95. Best matching sequences are enclosed in frames. The start codon for ORFs AC1 and AV1, the origin of replication (Ori), and the putative template exchange point that could promote the recombination event are indicated. The AC1 binding site and the conserved nona-nucleotide are highlighted. (**B**) Alignment of the IRs of DNA B1 and clone JJ95 (now called DNA A1). A ~140–nt–long common region is enclosed in frames. The AC1 binding site and the origin of replication are highlighted.

Since both *S. micrantha* and *M. parviflora* belong to the family Malvaceae and considering that DNA A2 accumulated predominantly (over DNA A1) in *M. parviflora* (see Fig. 6) and *S. micrantha* plants (Jovel *et al.*, 2004), it is likely that the symptomatic phenotype in the original *S. micrantha* plants in mainly induced by DNAs A2/B2. In contrast, studies with *Tomato mottle virus* (ToMoV) and *Bean dwarf mosaic virus* (BDMV), two begomoviruses isolated from a solanaceous and a leguminous species, respectively, showed that BDMV was more pathogenic in *N. benthamiana* and *N. tabacum* plants (two solanaceous species), than ToMoV (Hou *et al.*, 1998). Therefore, a conclusive resolution of the causal agent of the *Sida micrantha* mosaic will await until the Koch’s postulates could be fulfilled.

### Natural pseudo-recombinants

Several independent lines of evidence suggest that the AC1 protein of begomoviruses is a DNA-binding protein with nicking and closing activities and acts in a virus-specific manner (Hou and Gilbertson, 1996; Orozco *et al.*, 1998). The specificity is partially determined by a direct repeat preceding the TATA box for complementary transcription, which is identical between cognate components of a begomovirus but divergent in heterologous ones. The repeats match the sequence 5’–**GG–AG**TAYY**GG–AG** (Hanley-Bowdoin *et al.*, 1999). In the case of DNA A2 and DNA B2 of SimMV the common binding motif is **GGGG**taat**GGGG**. Pseudorecombination is a type of reassortment which occurs between two different begomoviruses that exchange their viral chromosomes (Unseld *et al.*, 2000b). In this paper, we showed that DNA A1 and DNA A2 of SimMV were able to transreplicate only DNAs B with identical binding sites. Attempts to create pseudorecombinants between DNA A of AbMV, SiGMCRV, SiGMVHV, or SiYVV, and any of the DNAs B of the SimMV, failed although several strategies were used (unpublished data). Comparing the sequences of the CR of AbMV, SiGMCRV, and SiYVV, a consensus sequence for the putative AC1 binding site is found (**GGAG**tatt**^A^_G_GAG**). These iterons are not compatible with the ones in SimMV, which explains the inability of these three DNAs A to trans-replicate any of the DNAs B of SimMV.

For *Tomato golden mosaic virus* (TGMV), mutational analysis revealed that both repeats are necessary for AC1 binding and origin recognition. The length of the spacer within and between repeats was essential for specific AC1 binding (Orozco *et al.*, 1998). Correspondingly, we found a natural pseudo-recombination event (plants 531 and 541), in which the AC1 binding site in DNA B2 (**GGGG**taat**GGGG**) could be recognized by an heterologous DNA A, presumably belonging to Macropthiliun mosaic geminivirus, possessing a slightly different AC1 binding site (**GGGG**aact**GGGG**), which matches the pre-requisites postulated by Orozco *et al*., (1998).

A second natural pseudorecombinant was found between SiYVV DNA A and SiGMCRV DNA B, which had already been obtained under experimental conditions (Unseld *et al.*, 2000a). These viruses possess compatible AC1 binding sites. It seems unlikely that *B. tabaci* selectively acquired SiYVV DNA A and SiGMCRV DNA B from the source plants and transmitted them to a single *M. parviflora* plant. A more plausible explanation is that both begomoviruses were transmitted to *M. parviflora* but SiYVV DNA A and SiGMCRV DNA B interacted in a more efficient way with host factors required for successful infection.

On the other hand, the fact that the IR of DNA A3 shows an apparently recombinant sequence between DNA A2 and DNA A of ToRMV-[Ube] probably suggests that this latter could have originated from a *S. micrantha-infecting* begomovirus. It is possible that DNA A3 of the SimMV complex is a parental form of ToRMV-[Ube] DNA A. Thus, evidence for both pseudo-recombination and molecular recombination was found in the analyses of the contaminated plants.

Collectively, our results demonstrate that pseudo-recombination and molecular recombination may take place under quasi-natural conditions when two or more begomoviruses coincide in the same infected plants. On the one hand, as suggested by the severe symptoms induced by the pseudo-recombinant SiYVV DNA A and SiGMCRV DNA B in plant 551 (Fig. 3B), and as has been found experimentally (Hou and Gilbertson, 1996), pseudo-recombination may give rise to recombinant viruses with increased virulence. Moreover, molecular recombination strongly promotes the diversity of begomoviruses. With this scenario in mind, strategies aimed at reducing the economical impact of geminiviral diseases in crop fields should be oriented towards the induction of plant defence mechanisms that are commonly exploited for several begomoviruses.

## Acknowledgements

We thank S. Kober and Diether Gotthardt for their skilful assistance. To the *Deutscher Akademischer Austausch Dienst* (DAAD) for financing the PhD studies of JJ and HJ.

## References

Abouzid, A. M., and Jeske, H. (1986). The purification and characterization of gemini particles from Abutilon mosaic virus infected Malvaceae. J. Phytopath. 115, 344–353.

Bedford, I. D., Briddon, R. W., Jones, P., Alkaff, N., and Markham, P. G. (1994). Differentiation of three whitefly– transmitted geminiviruses from the Republic of Yemen. Europ. J. Plant Pathology 100, 243–257.

Briddon, R. W., and Markham, P. G. (2001). Cotton leaf curl disease. Virus Res. 71, 151–159.

Czosnek, H., and Laterrot, H. (1997). A worldwide survey of tomato yellow leaf curl viruses. Arch. Virol. 142, 1391–1406.

Eigen, M. (1993). Viral quasispecies. Sci. Amer. 269(1), 32–39.

Frischmuth, T., Engel, M., Lauster, S., and Jeske, H. (1997). Nucleotide sequence evidence for the occurrence of three distinct whitefly-transmitted, Sida-infecting bipartite geminiviruses in Central America. J. Gen. Virol. 78, 2675–2682.

Hanley-Bowdoin, L., Settlage, S. B., Orozco, B. M., Nagar, S., and Robertson, D. (1999). Geminiviruses: Models for plant DNA replication, transcription, and cell cycle regulation. Crit. Rev. Plant Sci. 18, 71–106.

Höfer, P., Engel, M., Jeske, H., and Frischmuth, T. (1997a). Host range limitation of a pseudorecombinant virus produced by two distinct bipartite geminiviruses. Mol. Plant-Microbe Interact. 10, 1019–1022.

Höfer, P., Engel, M., Jeske, H., and Frischmuth, T. (1997b). Nucleotide sequence of a new bipartite geminivirus isolated from the common weed Sida rhombifolia in Costa Rica. J. Gen. Virol. 78, 1785–1790.

Höhnle, M., Höfer, P., Bedford, I.D., Briddon, R.W., Markham, P.G. and Frischmuth, T. (2001). Exchange of three amino acids in the coat protein results in efficient whitefly transmission of a nontransmissible Abutilon mosaic virus isolate. Virology 290(1), 164–71.

Hou, Y. M., and Gilbertson, R. L. (1996). Increased pathogenicity in a pseudorecombinant bipartite geminivirus correlates with intermolecular recombination. J. Virology 70, 5430–5436.

Hou, Y. M., Paplomatas, E. J., and L., G. R. (1998). Host adaptation and replication properties of two bipartite geminiviruses and their pseudorecombinants. Mol. Plant– Microbe Interact. 11, 208–217.

Jeske, H., Lütgemeier, M., and Preiss, W. (2001). Distinct DNA forms indicate rolling circle and recombination-dependent replication of Abutilon mosaic geminivirus. EMBO J. 20, 6158–6167.

Jovel, J., Preiss, W., Jeske, H. 2007. Characterization of DNA– intermediates of an arising geminivirus. Virus Res. In press.

Jovel, J., Reski, G., Rothenstein, D., Ringel, M., Frischmuth, T., and Jeske, H. (2003). Sida micrantha mosaic is associated with a complex of begomoviruses different from Abutilon mosaic virus. Arch. Virol. 149, 829–841.

Jovel, J. and Ruiz, E. 2004. Pathogenicity of the *Sida golden mosaic Costa Rica virus* (SiGMCRV) in several crop plants. Manejo Integrado de Plagas (Costa Rica) 70:19–29.

Morales, F. J., and Anderson, P. K. (2001). The emergence and dissemination of whitefly-transmitted geminiviruses in Latin America. Arch. Virol. 146(3), 415–441.

Orozco, B. M., Gladfelter, H. J., Settlage, S. B., Eagle, P. A., and Gentry, R. N. H.-B. L. (1998). Multiple cis elements contribute to geminivirus origin function. Virology 242(2), 346–356.

Pascal, E., Gooflove, P.E., Leeju, C.W., and Lazarowitz, S.G. (1993). Transgenic tobacco plants expressing the geminivirus BL1 protein exhibit symptoms of viral diseases. Plant Cell 5, 795–807.

Rybicki, E. P., Briddon, R. W., Brown, J. K., Fauquet, C. M., Maxwell, D. P., Harrison, B. D., Markham, P. G., Bisaro, D. M., Robinson, D., and Stanley, J. (2000). Family Geminiviridae. In “Virus Taxonomy – Classification and nomenclature of viruses” (M. H. V. Regenmortel van, C. M. Fauquet, and D. H. L. Bishop, Eds.), pp. 285–297. Academic Press, San Diego.

Thottappilly, G. (1992). Plant virus diseases of importance to African agriculture. J. Phytopathol. 134(265–288).

Unseld, S., Ringel, M., Höfer, P., Höhnle, M., Jeske, H., Bedford, I. D., Markham, P. G., and Frischmuth, T. (2000a). Host range and symptom variation of pseudorecombinant virus produced by two distinct bipartite geminiviruses. Arch. Virol. 145, 1449–1454.

Unseld, S., Ringel, M., Konrad, A., Lauster, S., and Frischmuth, T. (2000b). Virus-specific adaptations for the production of a pseudorecombinant virus formed by two distinct bipartite geminiviruses from Central America. Virology 274, 179–188.

Zhang, S. C., Wege, C., and Jeske, H. (2001). Movement proteins (BC1 and BV1) of Abutilon mosaic geminivirus are cotransported in and between cells of sink but not of source leaves as detected by green fluorescent protein tagging. Virology 290, 249–260.

